# Chemoreception of *Meloidogyne incognita* and *Caenorhabditis elegans* on botanical nematicidals

**DOI:** 10.1101/274092

**Authors:** Robert Sobkowiak, Natalia Bojarska, Emilia Krzyżaniak, Karolina Wągiel, Nikoletta Ntalli

## Abstract

Plant–parasitic nematodes cause serious damage to various agricultural crops worldwide, and their control necessitates environmentally safe measures. Plant secondary metabolites of botanical origin are tested here–in to study their effect in *Meloidogyne incognita* locomotion, being this an important factor affecting host inoculation inside the soil. We compare the effect to the respective behavioral responses of the model organism *Caenorhabditis elegans*. The tested botanical nematicidals, all reported of activity against *Meloidogyne* sp. in our previous works, belong to different chemical groups of small molecular weight molecules encompassing acids, alcohols, aldehydes and ketones. Specifically we report on the attractant or repellent properties of trans–anethole, (E,E)–2,4–decadienal, (E)–2–decenal, fostiazate, and 2–undecanone. The treatments for both nematode species were made at sublethal concentration levels, namely 1mM (<EC_50_), and the chemical control used for the experiment was the commercial nematicide fosthiazate and oxamyl. According to our results, trans–anethole, decenal, and oxamyl act as *C. elegans* attractants. 2–undecanone strongly attracts *M. incognita*. These findings can be of use in the development of nematicidal formulates, contributing to the disruption of nematode chemotaxis to root systems.

## Introduction

The plant parasitic nematodes are a major biotic stress factor, affecting numerous crop productions, and the root–knot nematodes (RKN: *Meloidogyne* spp.) is the main crop damaging species of Solanaceae worldwide. ^[1]^ Most of the synthetic plant protection products are now of the market (91/4141/EEU) due to their harsh environmental impact and to date research is focused in investigating new means of nematodes control. The plant secondary metabolites, gain the major part of the scientific attention to be studied as alternatives, eco–friendly nematode control substances and efficacy is evaluated in terms of second stage juveniles (J2s) paralysis or biological cycle arrest in host roots. ^[2, 3]^ Nonetheless and to the best of our knowledge there are only a few studies on the repellent or attractant properties of botanical substances against phytonematodes, although the pest orientation inside the soil, signifying host infection, is clearly defined by chemical signals. In fact, the infective nematode stage responds to plant signals originating from root exudates or sites of previous nematode penetration to find and recognize their hosts inside the soil. ^[4]^ In that frame, recently was analyzed the chemical composition of tomato root exudates to study the allopathic effect towards *Meloidogyne incognita*, yielding mainly esters and phenols. ^[5]^ Other substances that have been found of repellent and attractant properties towards RKN inside the soil are carbon dioxide ^[6, 7]^, tannic acids, flavonoids, glycoside, fatty acids and volatile organic molecules. ^[8, 9]^ Nematode chemoreception towards a nematicidal compound could help amegliorate efficacy if of attractant properties. Formulates composed of synergistic binary mixtures of nematicidals with nematode attractants could enhance even more efficacy levels under field conditions. Thus it is mandatory to unreveal both the nematicidal and the attractant or repellent properties of plant secondary metabolites, if to be used as active ingredients in a nematicidal IPM compatible formulate. Recently it has been reported that the sensory perception of *M. chitwoodi* and *M. hapla* juveniles was altered after exposure to seaweed products ^[10]^ and the effect was attributed to chemical properties of ingredient compounds. ^[11]^ Perry ^[11]^ has even provided a useful generalized framework to visualize attractants by classifying them as long–distance, short–distance and local attractants. Root–knot nematodes are said to be attracted to the root area by long–distance attractants. Short distance attractants attract the nematodes to the roots themselves while local attractants are responsible for orientation to the preferred invasion site by endoparasitic nematodes. In our previous works we demonstrated the activity of botanical aldehydes and ketones on RKN ^[12-15]^ and we attributed this activity to a potential inhibition of the vacuolar–type H+–ATPase (V–ATPase) in J2s. ^[16]^ Additionally we demonstrated the nematicidal activity of (E,E)–2,4–decadienal, 2–undecanone, furfural and (E)–2–decenal on J2 cuticle and eggs. ^[17]^

In this study, we report for the first time of the chemotaxis effects of trans–anethole, (E,E)–2,4– decadienal, (E)–2–decenal, fostiazate, 2–undecanone, fosthiazate, and oxamyl on *M. incognita* and the model nematode *C. elegans*.

## Materials and methods

### Nematodes rearing

#### Meloidogyne incognita maintenance

A pure population of *M. incognita* was reared from a single eggmass on tomato plants *Solanum lycopersicum* cv. Belladonna for two months. Plants used for inoculations were 7 weeks old at the five-leaf stage. All plants were maintained in a growth chamber at 25-28 °C, 60% relative humidity and 16 h photoperiod. After 60 days galled roots were separated from the soil, washed in tap water, and cut in pieces of 0.5 cm length. Eggs were extracted according to Hussey and Barker ^[18]^ those second-stage juveniles (J2) of *M. incognita* that hatched after 24–48 h were collected to be used in the experiments.

#### Caenorhabditis elegans maintenance

All tests were performed on the wild–type Bristol N2 strain of *C. elegans* obtained from the Caenorhabditis Genetics Center (CGC) at the University of Minnesota (Duluth, Minnesota, USA). Standard methods were used for the maintenance and manipulation of strains. ^[19, 20]^ Nematodes were kept at 22°C on nematode growth medium (NGM) agar plates seeded with *Escherichia coli* strain OP50 as a source of food according to a standard protocol. ^[21]^ Chunks of mixed–stage starved worms were transferred onto nematode growth medium (NGM) plates (50 mm in diameter) seeded with bacterial food (*E. coli* OP50), and the worms were allowed to grow for about 5 days at 22°C.

### Chemicals

(E,E)–2,4–decadienal, 2–undecanone, furfural and (E)–2–decenal were purchased from Sigma Aldrich.

### Behavioral assay

In all the experiments we used young adult worms exposed to 0 mM (control) and gradient (< 5 mM) of each of the test compounds. The single young adult hermaphrodites were transferred to assay plates containing NGM medium. The behavioral assays were performed in standard Petri dishes (inner diameter 92 mm) containing 5.8 mL of NGM. Then 1 μL of 5 mM test solution, prepared as explained in the above paragraph, was placed at the center of the plate just before the tracking. To study the behavior of worms, we used an automated tracking system to follow individual young adults crawling on NGM plates. ^[22^,^23]^ Single young adults (*C. elegans* aged ∼ 55h) or freshly hatched (24 h) *M. incognita* second–stage juveniles (J2) were manually picked off a plate and placed using a platinum pick in a 1 μL droplet of water on the agar surface of the gradient assay plate, 15 mm from the plate center (Fig. 1). Putting a worm in a droplet of water is an effective method for rapid transferring of a single animal without scratching the agar surface (important for obtaining high–contrast videos and perfect tracking) and for testing the condition of the worm (injured nematodes, which could not properly swim in a drop of water, were rejected). The surface tension of the drop of water prevented the nematode from creeping out. Next, the plate was placed in a device for tracking nematodes. Observation of the nematode allowed us to notice the moment when the 1 μL drop disappeared by evaporation/absorption in agar and released the worm, which could then freely move on the plate. At that moment, in the center of the plate, 1 μL of 5 mM test solution or 1 μL of carrier control (1% DMSO in water) was spotted, and the Petri dish was closed with a lid. After that, we recorded changes in their behavior in a substance gradient, where they could choose a desired substance concentration. Worm tracking began no more than 10 s after the application of test solutions or water in the center of the plate. Each worm was tracked for 3600 s or until it reached the edge of the plate.

**Figure 1.**
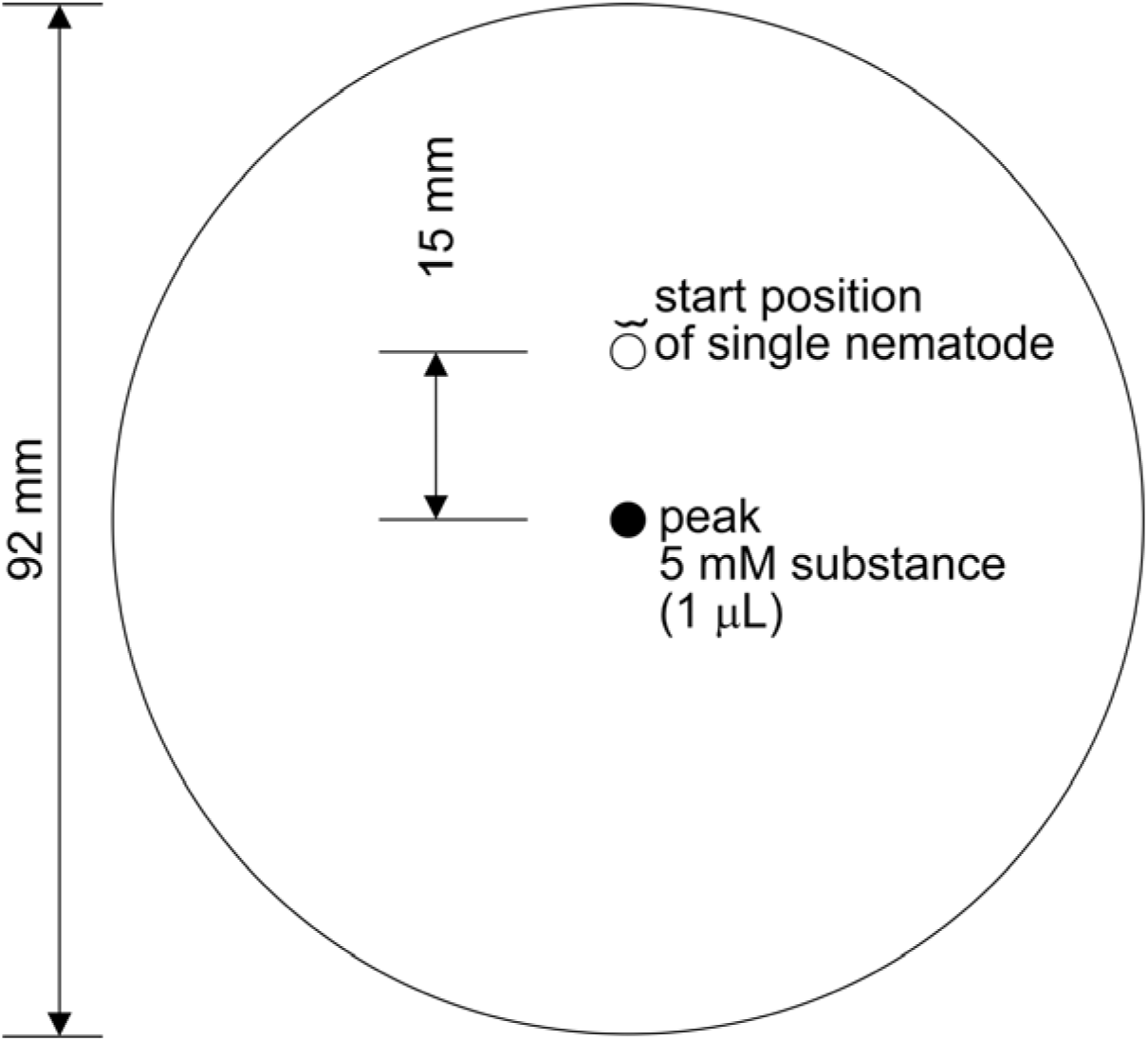
Configuration of the plate for substance gradient assay of nematode. One worm was placed in a 1 μL drop of water and thus trapped for several dozen seconds in the start position of a Petri dish containing 5.8 mL of NGM medium. Immediately after the worm was released, substance was spotted into the center of the plate (peak). Water or 2% DMSO was applied instead of substance in control group.

#### Worm tracking system

Custom worm tracker software (WormSpy) was used to move the camera automatically to re–center the worm under the field of view during recording. ^[22^,^23]^ The automated tracking system comprises a stereomicroscope (Olympus SZ11), a modified (with unscrewed lens) web camera (Logitech QuickCam Pro 9000) with 640×480 resolution to acquire worm videos, and a desktop PC running under Windows 7. The tracking system located the worm’s centroid (defined as the geometrical center of the smallest rectangle that could be drawn around the worm) and recorded its x and y coordinates with a sampling rate of 2 s^-1^. When a worm neared the edge of the field of view, the tracking system automatically re–centered the worm by moving the stage and recorded the distance that the stage was moved. We reduced the variation in sampling rate as a consequence of the small differences in the time it took to re–center the worm and the need to take data only when the stage was stationary by developing a simultaneous localization and tracking method for a worm tracking system. ^[22^,^23]^ The spatiotemporal track of each worm was reconstructed from the record of centroid locations and camera displacements. The instantaneous speed and trajectory were computed using the displacement of the centroid in successive samples. The tracking system recorded the worm’s position, speed, and distance from the center of the plate and from the starting point, trajectory at 0.5–s intervals. Individual worms moved away from their starting location, leaving complex tracks. Video recordings were carried out at room temperature (22 °C). Thirty independent experiments were performed per group. All the experimental procedures presented in this paper were in compliance with the European Communities Council Directive of 24 November 1986 (86/609/EEC).

### Statistical Analysis

The measurements from experiments were pooled for each treatment group and the median values were calculated. The data were not normally distributed, as determined by the Shapiro–Wilk W–test and Kolmogorov–Smirnov & Lilliefors method. Due to this, the Kruskal–Wallis tests followed by Dunn’s multiple comparison post–hoc tests were performed. Statistical significance was considered at p < 0.001. The calculations and graphs were done by using Statistica software (StatSoft, Inc., Tulsa, Oklahoma, USA).

## Results

### Average values of factors describing C. elegans and M. incognita behavior

#### Average speed of movement

The average speed of movement of *C. elegans* worms (Fig. 2A) was distinctly faster to the respective control in presence of (E,E)–2,4–decadienal. The average speed of movement *C. elegans* worms (Fig. 2A) was slower to DMSO in the descending order of (E)–2–decenal followed by (E,E)–2,4–decadienal and trans–anethol. Oxamyl was slightly slower to water and fosthiazate slightly faster. The average speed of movement *M. incognita* worms (Fig. 6A) was distinctly faster in presence of (E)–2–decenal in comparison to DMSO.

**Figure 2.**
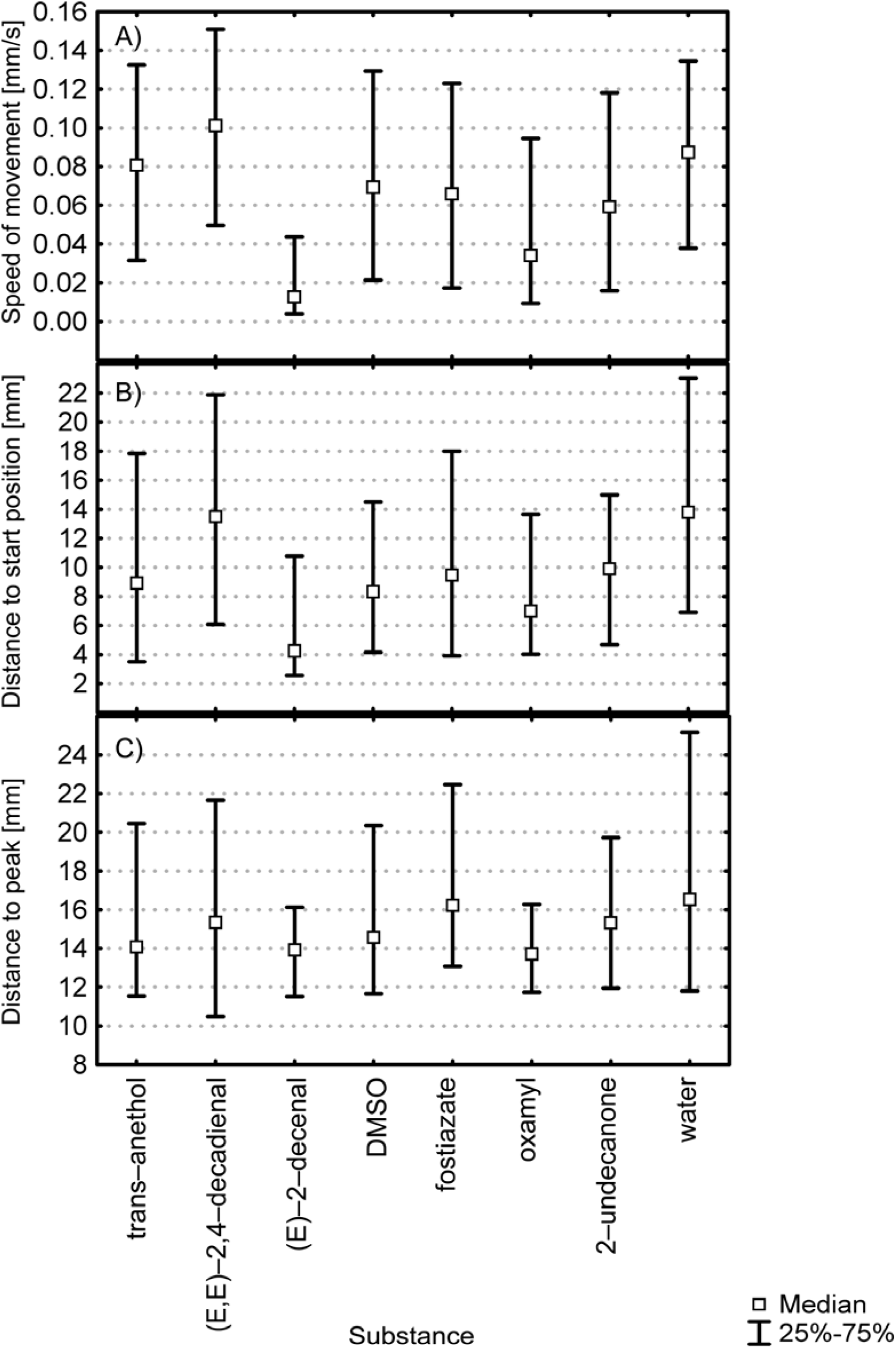
Average values of factors describing the behavior of *Caenorhabditis elegans*: (A) speed of movement; (B) distance from the start position; (C) distance from the peak; All groups were statistically significantly different from each other (p < 0.001, Kruskal–Wallis ANOVA by ranks test, data pooled from 30 independent experiments, N > 45 400).

#### Average distance from the start position

*C. elegans* moved away from the starting point in a range of 4 mm ((E)–2–decenal) to 14 mm (water) (Fig. 2B). *M. incognita* moved away from the starting point in a range of 3 mm (trans– anethol and water) to 4.5 mm (2–undecanone) (Fig. 6B).

Some substances can quickly cause paralysis or quiescence (some kind of sleeping). Please see case *C. elegans* and (E)–2–decenal (Fig. 2A lowest speed of movement, Fig. 2B lowest distance to start position). This could mean that worm is paralysed/anaesthetized.

#### Average distance from the peak

The peak was the central point where substance or control solution was applied on the gradient assay plate. The radius in which *C. elegans* were present, measured from the peak to worm location, was in the range from about 14 mm (trans–anethol, (E)–2–decenal, oxamyl) to about 16 mm (water and fostiazate) (median value, Fig. 2C).

The radius in which *M. incognita* were present, measured from the peak to worm location, was in the range from about 14.5 mm ((E)–2–decenal) to about 11.5 mm (2–undecanone) (median value, Fig. 6C). (E)–2–decenal is repellent (Fig 5C). 2–undecanone is strong attractant for *M. incognita* (Fig 5C).

### Time–dependent response of C. elegans and M. incognita to the test compounds’ gradient

#### Time–dependent speed of movement

The speed of the *C. elegans* varied with time (Fig. 3). In most cases, we observed a decreasing trend in the speed of its movements, and its frequent oscillations. In control conditions (Fig 2A and B), we observed lowest oscillations of the speed, while the nematodes in presence of (E)–2–decenal were paralysed (Fig. 6E). The long period of reduced locomotion activity was observed particularly in presence of oxamyl and 2–undecanone (Fig. 3G and H, time 1800–3400 s).

**Figure 3.**
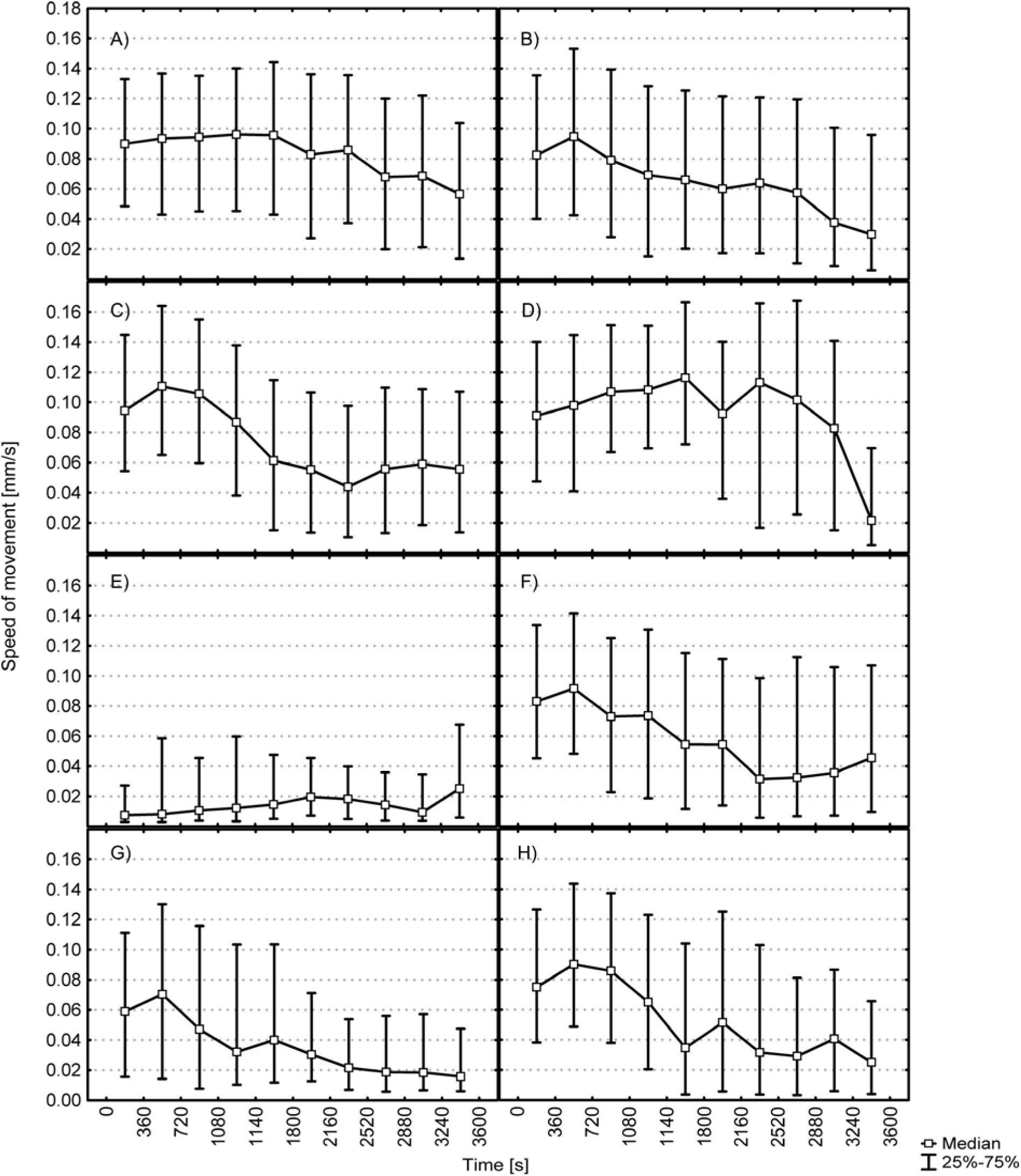
Locomotor activity (centroid speed) of *Caenorhabditis elegans* in the tested experimental variants. The data are medians and 25th and 75th percentiles of 30 pooled experiments. (A) water; (B) DMSO; (C) trans–anethol; (D) (E,E)–2,4–decadienal; (E) (E)–2–decenal; (F) fostiazate; (G) oxamyl; (H) 2–undecanone.

The speed of the *M. incognita* was more stable in comparison to *C. elegans* (Fig. 7). In most cases, we observed a more or less side trend in the speed of their movements. That is *M. incognita* moves with sustain almost stable speed themselves over the long time. The oscillations were revealed in control conditions (water Fig. 7A). The long period of reduced locomotion activity was particularly observed in presence of fostiazate (Fig. 7F).

#### Time–dependent changes in distance from the start position

Figure 4 shows the variation in the distance from the start position of *C. elegans* (measured from the start position to current worm location). Generally during the experiment the nematodes moved away from the starting point. Some substances can quickly cause paralysis or quiescence (some kind of sleeping). Please see case *C. elegans* and (E)–2–decenal (Fig. 2A lowest speed of movement, Fig. 2B lowest distance to start position this could mean that worms sleep). In some experimental conditions, nematodes remained at a relatively stable distance from the start position in presence of (E)–2–decenal (Fig. 4E) and oxamyl (Fig. 4G). The greatest distance from the start position was recorded for the nematodes in presence (E,E)–2,4–decadienal (Fig. 4D).

**Figure 4.**
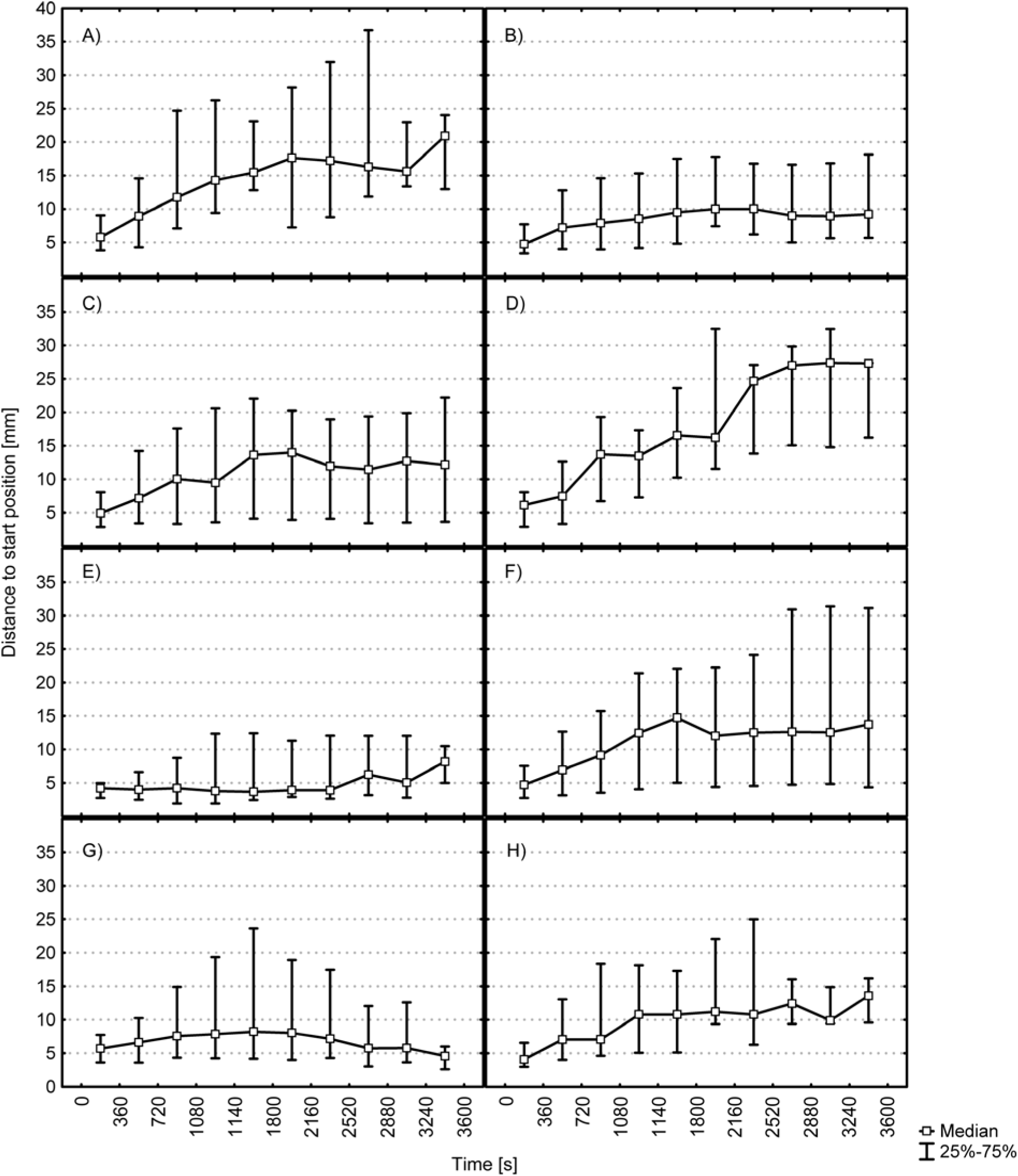
Distance from the start position of *Caenorhabditis elegans* in the tested experimental variants. The data are medians and 25th and 75th percentiles of 30 pooled experiments. (A) water; (B) DMSO; (C) trans–anethol; (D) (E,E)–2,4–decadienal; (E) (E)–2–decenal; (F) fostiazate; (G) oxamyl; (H) 2–undecanone.

**Figure 5.**
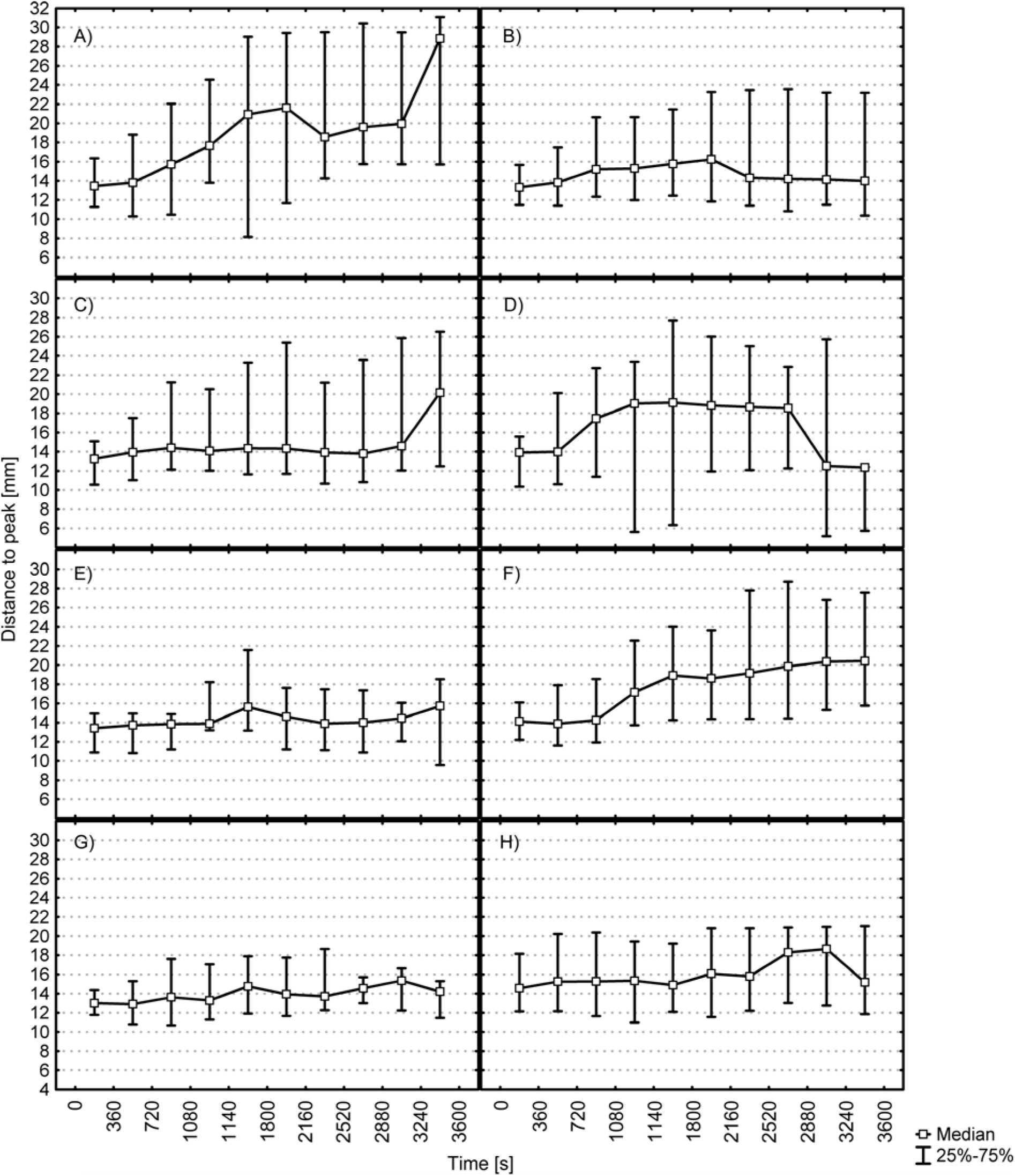
Time–dependent changes in distance from the peak of *Caenorhabditis elegans* in the tested experimental variants. The data are medians and 25th and 75th percentiles of 30 pooled experiments. (A) water; (B) DMSO; (C) trans–anethol; (D) (E,E)–2,4–decadienal; (E) (E)–2– decenal; (F) fostiazate; (G) oxamyl; (H) 2–undecanone.

**Figure 6.**
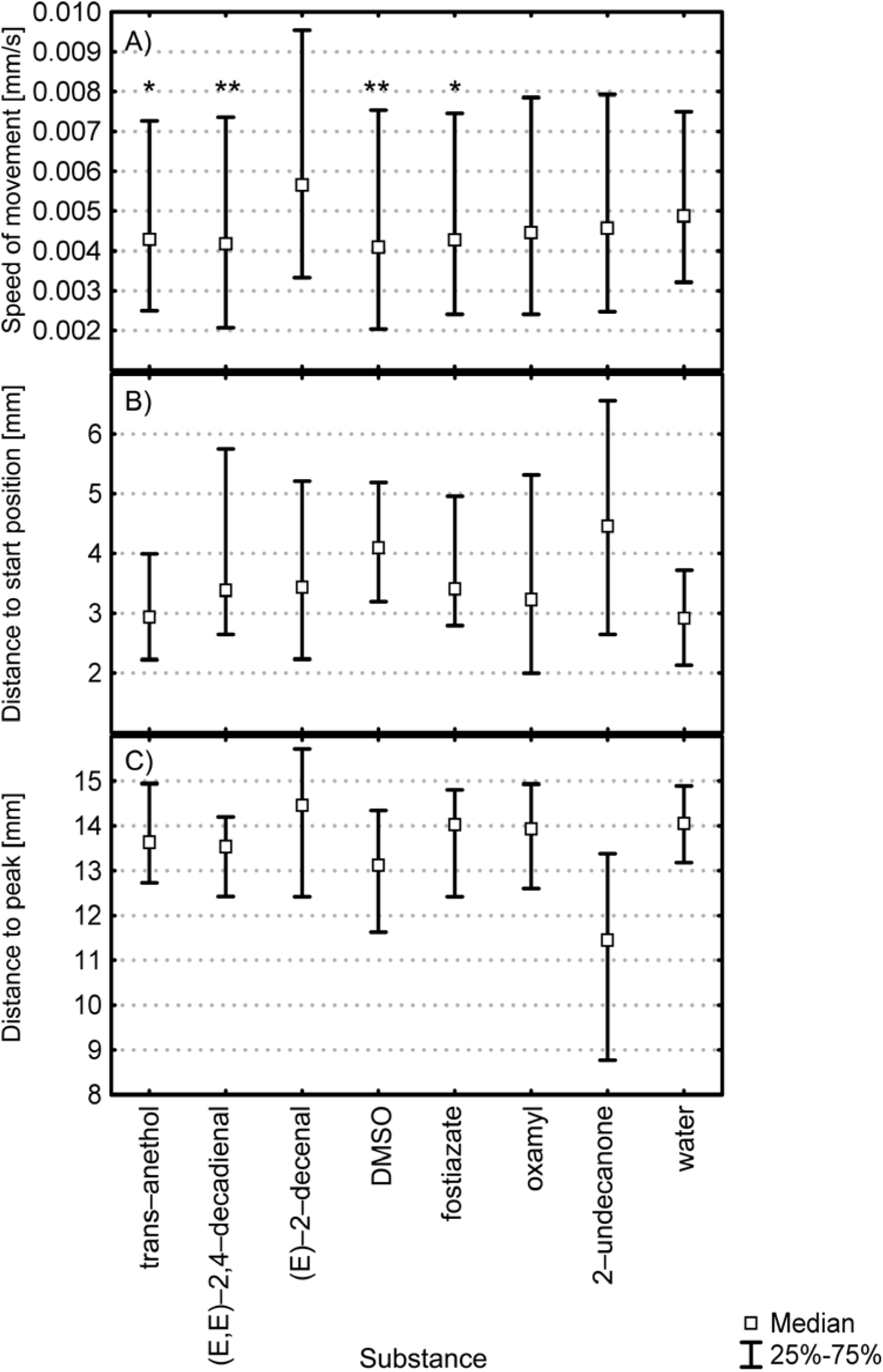
Average values of factors describing the behavior of *Meloidogyne incognita*: (A) speed of movement, Fostiazate vs. trans–anethol (*) and DMSO vs. (E,E)–2,4–decadienal (**) were not statistically significantly different; (B) distance from the start position; (C) distance from the peak. Other groups were statistically significantly different from each other (p < 0.001, Kruskal–Wallis ANOVA by ranks test, data pooled from 30 independent experiments, N > 65 000).

Figure 8 shows the variation in the distance from the start position of *M. incognita*. During the experiment the nematodes moved away from the starting point with exception when on the center of the Petri dish was applicated water (Fig. 8A) or DMSO (Fig. 8B). Farthest median distance from the start position was recorded for *M. incognita* in presence (E)–2–decenal (Fig. 8E).

**Figure 7.**
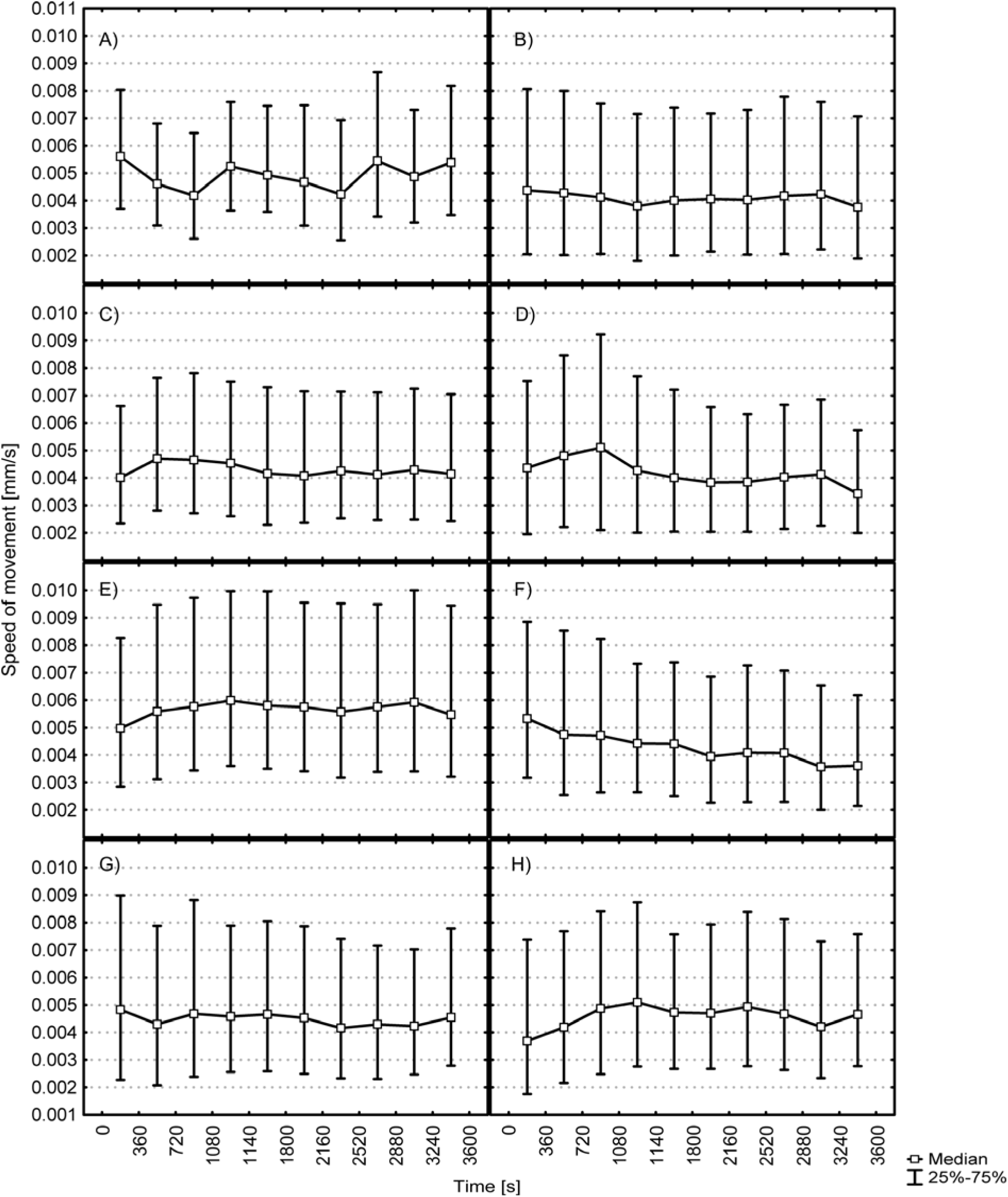
Locomotor activity (centroid speed) of *Meloidogyne incognita* in the tested experimental variants. The data are medians and 25th and 75th percentiles of 30 pooled experiments. (A) water; (B) DMSO; (C) trans–anethol; (D) (E,E)–2,4–decadienal; (E) (E)–2–decenal; (F) fostiazate; (G) oxamyl; (H) 2–undecanone.

**Figure 8.**
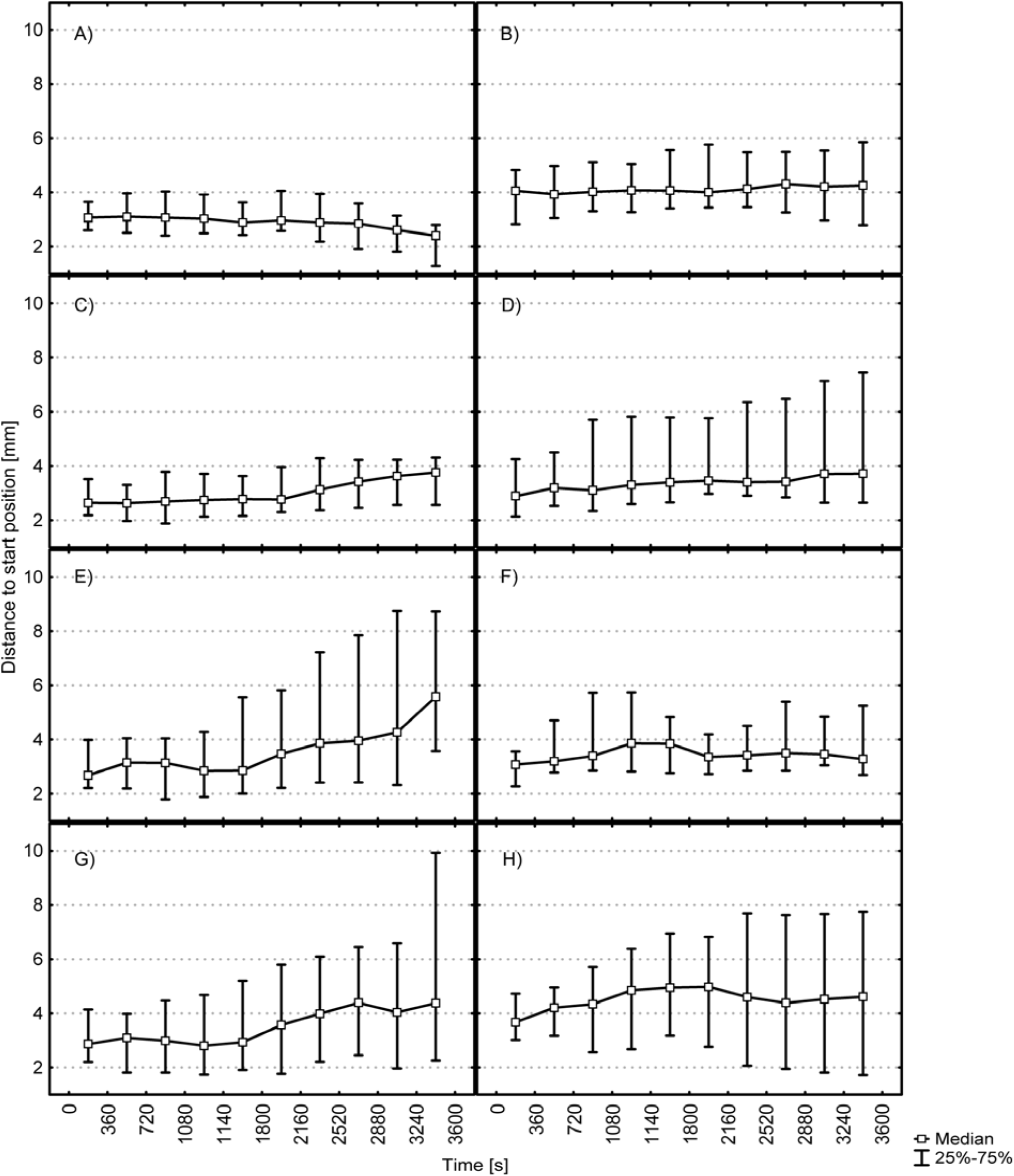
Distance from the start position of *Meloidogyne incognita* in the tested experimental variants. The data are medians and 25th and 75th percentiles of 30 pooled experiments. (A) water; (B) DMSO; (C) trans–anethol; (D) (E,E)–2,4–decadienal; (E) (E)–2–decenal; (F) fostiazate; (G) oxamyl; (H) 2–undecanone.

#### Time–dependent changes in distance from the peak

The distance between *C. elegans* and the peak varied with time (Fig. 5). The worms generally did not approach to the center of plate. Most often *C. elegans* is keen to escape. Only little approach was observed in presence of (E,E)–2,4–decadienal (Fig. 5D, A)

Interesting behavior of *M. incognita* was observed in presence of 2–undecanone where almost immediately, in time range of 0 – 360 s, after application nematodes moved towards the central peak of the test substance (Fig. 9H). Fostiazate evoked moving towards the central peak only about half duration of experiments (Fig. 9F).

**Figure 9.**
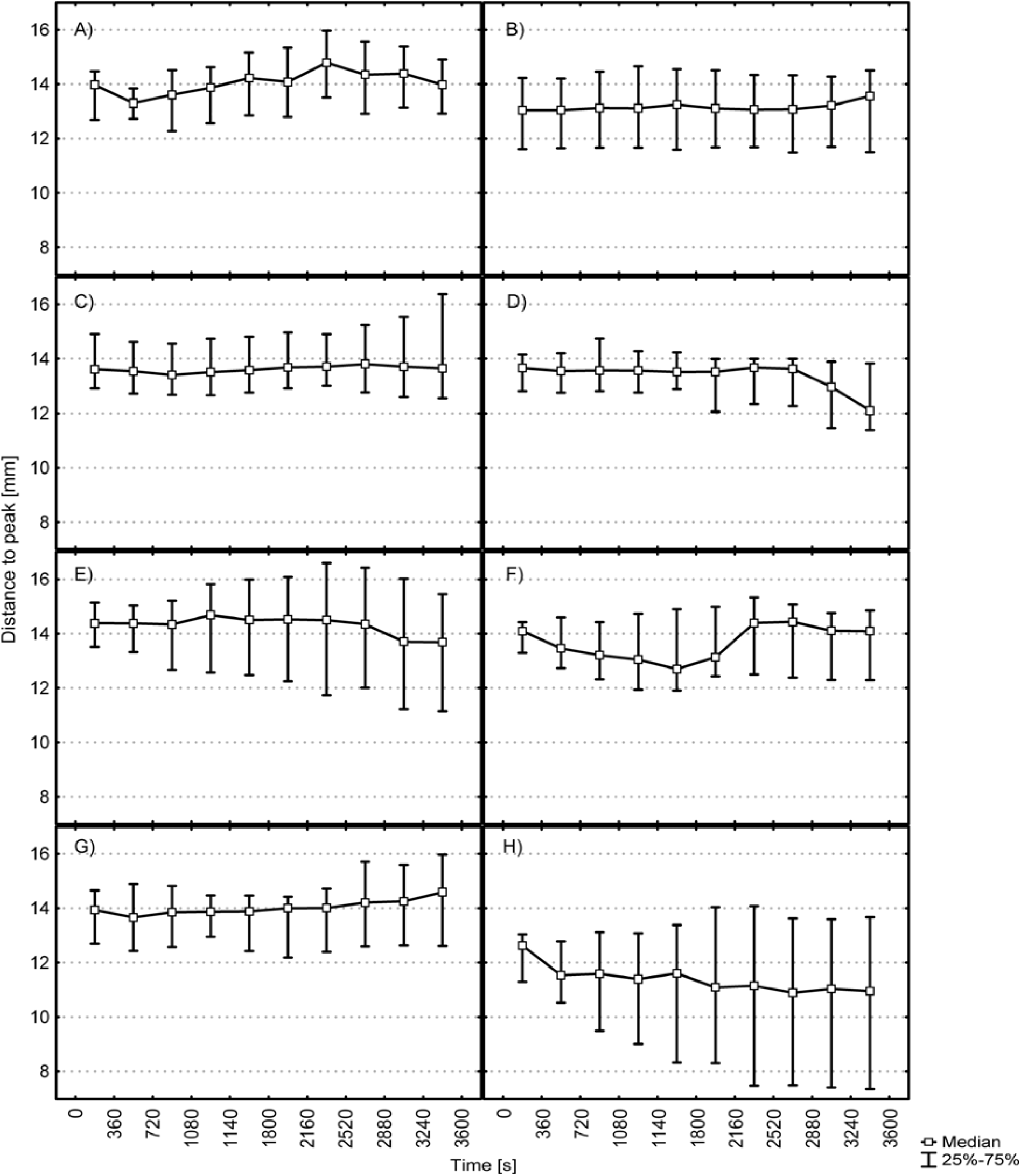
Time–dependent changes in distance from the peak of *Meloidogyne incognita* in the tested experimental variants. The data are medians and 25th and 75th percentiles of 30 pooled experiments. (A) water; (B) DMSO; (C) trans–anethol; (D) (E,E)–2,4–decadienal; (E) (E)–2– decenal; (F) fostiazate; (G) oxamyl; (H) 2–undecanone.

## Discussion

Among plant–parasitic nematodes, root–knot nematodes (RKN), *Meloidogyne* spp., including *M. incognita*, are the most important provoking billions of euros of crop losses annually. Besides the plant parasitic forms, there are a large group of saprophytic nematodes found in the soil among which *C. elegans*. Laboratory experiments and field studies have demonstrated that saprophytic nematodes like *C. elegans* play a critical role in influencing the turnover of the soil microbial biomass and thus enhance the availability of plant nutrients. ^[24]^ Additionally there is an increasing interest in the use of *C. elegans* as a tool for parasitic nematode research. ^[25]^ In some ecosystems nematodes contribute up to 40% of nutrient mineralization. ^[26]^ The soil nematodes like *Caenorhabditis elegans*, also mediate the interaction between roots and rhizobia in a positive way, leading to nodulation. ^[27]^ Plant roots trigger behavioral response by emitting volatile signals attracting parasitic nematodes to the root proximity. ^[27]^

In order to increase agricultural production yields the plant protection products are used. Commercial nematicides may effectively control nematode pests in annual crops, but are usually non–specific, notoriously toxic and they pose a threat to saprophytic nematodes, and the soil ecosystem, ground water and human health. ^[28]^

During the past decade, the roundworm *Meloidogyne incognita* has become a popular target for the study of nematicides, leading to a wealth of information about toxic substances of plant origin ^[2]^ in order to replace biohazardous nematicides with ecofriendly alternatives. Trans–anethole, (E,E)–2,4– decadienal, (E)–2–decenal, 2–undecanoneare are reported nematicidals ^[13^-^15]^ but their chemotactic effect on *M. incognita* J2 stage and adult *C. elegans* behavior is presented herein for the first time. The variability of the distribution of the nematicide in soil depends upon the degree of incorporation and the extent to which the chemical is redistributed in soil by diffusion and leaching in soil water.^[29]^ Failure to effectively control nematodes may often be explained by poor nematicide application and/or incorporation with field equipment, and the fact that the nematodes would move away from a repellent source when are exposed to higher concentrations, because the nematodes have developed the capacity to sense and respond to chemical signals. ^[29]^ The development of new classes of nematicides with novel mode of action, that are effective when used in soil or applied directly to crops and are environmentally benign and specific to target pests, is perhaps an idealistic hope. To be effective, nematicides have to persist long enough for nematodes to be exposed to lethal concentrations. One solution to this problem could be searching substances specifically attracting *M. incognita*.

We analyzed the movement of *M. incognita* J2 larvae and adult *C. elegans* in the presence of different concentrations of commercial nematicides (oxamyl, fostiazate) and plant derived compounds (trans–anethol, (E,E)–2,4–decadienal, (E)–2–decenal, and 2–undecanone) nematicides to investigate their chemotaxis and infer their responses to chemosensation *in vivo*. In our experiment, nematodes could select the concentration of substance in which they wanted to stay. Chemical components may deter one organism while attracting another and these compounds alter nematode behavior and can either attract nematodes or result in repellence, motility inhibition with narcotic effect or even death. ^[4]^ We used sublethal nematocides concentrations, but high enough to be “perceived” by the nematodes. One line of defense against the harmful substances is behavior; *C. elegans* and *M. incognita* can sense the presence of certain chemicals and avoid them.

In a survey of volatile organic compounds, *C. elegans* exhibited either attraction or repulsion to 50 out of 120 compounds tested. ^[30]^ Root exudates effect on chemotaxis of *M. incognita* J2. ^[5]^ Unlike free–living nematodes such as *C. elegans*, which feed on a wide range of bacterial species as well as fungal mycelium, parasitic nematodes must fine–tune their chemosensory repertoire to respond more precisely to host–specific cues. Because of different biology, the some substances evoke different behavior in *C. elegans* and *M. incognita*. Among tested substances the strongest repellent for *M. incognita* was (E)–2–decenal (Fig. 6C, 7E). 2–undecanone was strong attractant for *M. incognita* and the worms moved without any delay towards the substance (Fig. 6C, 8H). So far has been shown that carbon dioxide released from roots attracts *M. incognita.* ^[7]^ Yang et al. ^[5]^ revealed that sterilized water (control) attracted *M. incognita* J2. Likewise, in our experiment in water control *M. incognita* stay nearest to start position (Fig. 6B, 7A).

All tested compounds paralysed *M. incognita* and (E,E)–2,4–decadienal was the most effective followed by trans–anethole, (E)–2–decenal and 2–undecanone. ^[12^-^15]^ As 2–undecanone is the only attractant it would be interesting to study its efficacy in arresting the biological cycle of *M. incognita* in tomato roots and compare it to the repellents efficacy.

Paralysis we observed when *C. elegans* were challenged to (E)–2–decenal (see speed of movement and distance to start position Fig. 2, 3E, 4E). The similar narcotic effect we observed in presence of oxamyl – commercial non fumigant nematicide which acts as acetylcholinesterase inhibitors, interfering with the normal nerve impulse transmission within the central nervous system of nematodes.

## Conclusions

To the best of our knowledge, this is the first report on the behavioral effects of trans–anethole, (E,E)–2,4–decadienal, (E)–2–decenal, 2–undecanone are, fosthiazate and oxamyl on *Meloidogyne incognita* and *Caenorhabditis elegans*. Moreover, the data on the behavior and neural activity in response to spatial chemical patterns of tested substances are too limited to make a definitive assessment. Therefore, further studies on the suborganismal level can explain the mode of toxic activity of aldehydes and ketones in cells and tissues. ^[17]^ Our results suggest that natural substances may significantly decrease crop loss due to changed nematode behavior. Therefore, we think that even when lethality of natural substances is lower than lethality of chemical nematicides, they can like e.g. 2–undecanone acting synergistically to enhance attraction to synthetic pesticides.

## Acknowledgments

We thank W.R. Schafer and E. Yemini for discussion. We would like to thank the Caenorhabditis Genetics Center, which is funded by the NIH National Center for Research Resources (NCRR), for supplying the nematode strain used in this research. Special thanks to Tzortzakakis Emmanuel, Researcher Nematologist at N.AG.RE.F, Plant Protection Institute, Heraklion, Crete, Greece for kindly providing pure nematode species. This study was supported by funds from the Department of Cell Biology, Faculty of Biology, Adam Mickiewicz University Poznań, Poland and Benaki Phytopathological Institute, Department of Pesticides Control & Phytopharmacy, Laboratory of Biological Control of Pesticides, 8 Stefanou Delta Street, Kifissia, Athens, 14561.

## References

[1] Barbary, A.; Djian-Caporalino, C.; Palloix, A.; Castagnone-Sereno, P. Host genetic resistance to root-knot nematodes, *Meloidogyne* spp., in Solanaceae: from genes to the field. Pest. Manag. Sci. 2015, 71, 1591–1598.

[2] Ntalli, N.G.; Caboni, P. Botanical nematicides: a review. Agric. Food Chem. 2012, 60, 9929–9940.

[3] Ntalli, N.G.; Caboni, P. Botanical nematicides in the mediterranean basin. Phytochem. Rev. 2012, 11, 351–359.

[4] Curtis, R.H. Plant-nematode interactions: environmental signals detected by the nematode’s chemosensory organs control changes in the surface cuticle and behaviour. Parasite. 2008, 15, 310–316.

[5] Yang, G.; Zhou, B.; Zhang, X.; Zhang, Z.; Wu, Y.; Zhang, Y.; Lu, S.; Zou, Q.; Gao, Y.; Teng, L. Effects of tomato root exudates on *Meloidogyne incognita*. PLoS One. 2016, 11, e0154675.

[6] Dusenbery, D.B. Theoretical range over which bacteria and nematodes locate plant roots using carbon dioxide. J Chem. Ecol. 1987, 13, 1617–1624.

[7] Pline, M.; Dusenbery, D.B. Responses of plant-parasitic nematode *Meloidogyne incognita* to carbon dioxide determined by video camera-computer tracking. J. Chem. Ecol. 1987, 13, 873–888.

[8] Chitwood, D.J. Phytochemical based strategies for nematode control. Annu. Rev. Phytopathol. 2002, 40, 221–249.

[9] Rasmann, S.; Hiltpold, I.; Ali, J. The role of root–produced volatile secondary metabolites in mediating soil interactions. In Advances in lSelected Plant Physiology Aspects, Montanaro, G. Ed., InTech Open Access Publisher, 2012; 269–290.

[10] Massa Ngala, B.; Valdes, Y.; dos Santos, G.; Perry, R.N.; Wesemael, W.M.L. Seaweed–based products from *Ecklonia maxima* and *Ascophyllum nodosum* as control agents for the root–knot nematodes *Meloidogyne chitwoodi* and *Meloidogyne hapla* on tomato plants. Appl. Phycol. 2016, 28, 2073–2082.

[11] Perry, R.N. An evaluation of types of attractants enabling plant-parasitic nematodes to locate plant roots. Russ. J. Nematol. 2005, 13, 83–88.

[12] Caboni, P.,; Aissani, N.; Cabras, T.; Falqui, A.; Marotta, R.; Liori, B.; Ntalli, N.; Sarais, G.; Sasanelli, N.; Tocco, G. Potent nematicidal activity of phthalaldehyde, salicylaldehyde, and cinnamic aldehyde against *Meloidogyne incognita*. J. Agric. Food Chem. 2013, 61, 1794–1803.

[13] Caboni, P.; Ntalli, N.G.; Aissani, N.; Cavoski, I.; Angioni, A. Nematicidal activity of (E,E)-2,4-decadienal and (E)-2-decenal from *Ailanthus altissima* against *Meloidogyne javanica*. J. Agric. Food Chem. 2012, 60, 1146–1151.

[14] Ntalli, N.G.; Vargiu, S.;Menkissoglu-Spiroudi, U.; Caboni, P. Nematicidal carboxylic acids and aldehydes from *Melia azedarach* fruits. J. Agric. Food Chem. 2010, 58, 11390–11394.

[15] Ntalli, N.G.; Manconi, F.; Leonti, M.; Maxia, A.; Caboni, P. Aliphatic ketones from *Ruta chalepensis* (Rutaceae) induce paralysis on root knot nematodes. J. Agric. Food Chem. 2011, 59, 7098–7103.

[16] Caboni, P.; Tronci, L.; Liori, B.; Tocco, G.; Sasanelli, N.; Diana, A. Tulipaline A: structure-activity aspects as a nematicide and V-ATPase inhibitor. Pestic. Biochem. Physiol. 2014, 112, 33–39.

[17] Ntalli, N.;Ratajczak, M.; Oplos, C.; Menkissoglu-Spiroudi, U.; Adamski, Z. Acetic acid, 2-undecanone, and (E)-2-decenal ultrastructural malformations on *Meloidogyne incognita*. Nematol. 2016, 48, 248–260.

[18] Hussey, R.S.; Barker, K.R. A comparison of methods of collecting inocula of *Meloidogyne* spp., including a new technique. Plant Dis. Rep. 1973, 57, 1025–1028.

[19] Stiernagle, T. Maintenance of *C. elegans*. WormBook, 2006, 1–11.

[20] Sobkowiak, R., Kowalski, M.; Lesicki, A. Concentration-and time-dependent behavioral changes in *Caenorhabditis elegans* after exposure to nicotine. Pharmacol. Biochem. Behav. 2011, 99, 365–370.

[21] Brenner, S. The genetics of *Caenorhabditis elegans*. Genetics, 1974, 77, 71–94.

[22] Kowalski, M.; Kaczmarek, P.; Kabaciński, R.; Matuszczak, M.; Tranbowicz, K.; Sobkowiak, R. A simultaneous localization and tracking method for a worm tracking system. International J.Appl. Math. Comp. Sci. 2014, 24, 599–609.

[23] Sobkowiak, R., Kaczmarek, P., Kowalski, M., Kabacinski, R., Lesicki, A. Behavior of *Caenorhabditis elegans* in a nicotine gradient modulated by food. Drug. Chem. Toxicol. 2017, 10.1080/01480545.2017.1405971.

[24] Bardgett, R.D.; Cook, R.; Yeates, G.W.; Denton, C.S. The influence of nematodes on below-ground processes in grassland ecosystems. Plant and Soil, 1999, 212, 23–33.

[25] Weaver, K.J.; May, C.J.; Ellis, B.L. Using a health-rating system to evaluate the usefulness of *Caenorhabditis elegans* as a model for anthelmintic study. PLoS One. 2017, 12, e0179376.

[26] De Ruiter, P.C.; Van Veen, J.A.; Moore, J.C.; Brussaard, L.; Hunt, H.W. Calculation of nitrogen mineralization in soil food webs. Plant and Soil, 1993, 157, 263–273.

[27] Horiuchi, J.; Prithiviraj, B.; Bais, H.P.; Kimball, B.A.; Vivanco, J.M. Soil nematodes mediate positive interactions between legume plants and rhizobium bacteria. Planta, 2005, 222, 848–857.

[28] Ntalli, N.; Monokrousos, N.; Rumbos, C.; Kontea, D.; Zioga, D.; Argyropoulou, M.D.; Menkissoglu-Spiroudi, U.; Tsiropoulos, N.G. Greenhouse biofumigation with Melia azedarach controls Meloidogyne spp. and enhances soil biological activity. J. Pest Sci. 2017, DOI 10.1007/s10340-017-0909-1.

[29] Mueller, J.D.; Noling, J.W. Standardization of reporting procedures for nematicide efficacy testing: a research and extension perspective. J. Nematol. 1996, 28, 575–585.

[30] Bargmann, C.I.; Hartwieg, E.; Horvitz, H.R. Odorant-selective genes and neurons mediate olfaction in *C. elegans*. Cell, 1993, 74, 515–527.

